# Calcium signals natural transformation in *Acinetobacter baumannii*

**DOI:** 10.64898/2026.02.23.707608

**Authors:** Jason Baby Chirakadavil, Kelly Goldlust, Ludovic Poiré, Nicolas Gaudin, Alexandre Chassard, Adrien Camilli, Manon Bouvier, Maria-Halima Laaberki, Xavier Charpentier

## Abstract

The acquisition of resistance to antibiotics in the opportunistic pathogen *Acinetobacter baumannii* may be linked to its capacity to undergo natural transformation. This mode of horizontal gene transfer relies on the import of extracellular DNA and its chromosomal integration by homologous recombination. Type IV pilus activity initiates the capture of extracellular DNA, which is then transported to the cytoplasm for recombination in the chromosome. While most *Acinetobacter baumannii* strains are transformable, the conditions allowing the expression of type IV pilus and other transformation genes remain largely unexplored. By investigating transformation-permissive conditions, we uncovered that calcium is a potent inducer of natural transformation. Type IV pilus genes and other transformation-specific genes (*comEA*, *dprA*) are upregulated by submillimolar concentrations of calcium ions, in a growth phase-dependent manner. In contrast, sodium chloride represses expression of *pilA*, counteracting the calcium-dependent induction, explaining the reported absence of transformation in NaCl-containing medium (such as LB). Independently of transcriptional induction, calcium ions also directly bind the type IV pilus through the calcium-dependent adhesin PilY1. Our data support a model in which calcium strengthens the interaction of PilY1 with the minor pilin complex, increasing pilus dynamics and subsequent pilus-dependent DNA capture. Hence, calcium signals natural transformation through both transcriptional and structural activation of type IV pilus. In addition to providing new insights into the regulation of natural transformation in *A. baumannii* this work led us to establish a protocol for genetic engineering of *A. baumannii* by natural transformation.

**Importance:** *Acinetobacter baumannii* is a nosocomial pathogen considered a critical research priority due to its resistance to last resort antibiotics. Understanding how *A. baumannii* evolves and acquires resistance to antibiotics is thus of prime importance. This species is capable of natural transformation, a means to acquire and spread genetic information, including antibiotic resistance genes. However, the conditions under which this process is active in this species remain elusive. We identify calcium ions as potent inducers of natural transformation and propose a model of the signaling of natural transformation by calcium ions. This opens the way for further investigations into the contribution of natural transformation to acquisition of antibiotic resistance. In addition, it provides an efficient way to genetically manipulate most *A. baumannii* strains.

## Introduction

Natural transformation is one of the main mechanisms of horizontal gene transfer driving the evolution of bacterial genomes (1). Over 80 bacterial species have been experimentally demonstrated to be capable of natural transformation (2). This ever-growing list includes *Acinetobacter baumannii*, a critical priority for research and development of new therapeutic strategies (3) due to its resistance to last resort antibiotics. Natural transformation likely plays a major role in the evolution of *A. baumannii* genomes, constituting a route of gene exchange promoting recombination events and acquisition of antibiotic resistance (4–6). Although natural transformation in *A. baumannii* appears to play a limited role in overall gene acquisition, at least 15% of antibiotic resistance genes had been acquired through this route (5). Indeed, even large genetic elements, such as AbaR resistance islands, could be acquired by natural transformation during the co-culture of *A. baumannii* isolates (7). Yet, it is the subsequent partial inhibition of natural transformation by these resistance islands that favors the prevalence of antibiotic resistance genes (8). Most *A. baumannii* are naturally transformable, but at varying levels (9–12). Such variations in natural transformation could be ascribed to genomic elements like restriction-modification systems that can limit transformation (11) and prophages inhibiting transformability (10). It remained possible that not all strains respond to the same stimuli for expression of transformation, yet little is known about the extracellular conditions that impact the expression of natural transformation in *A. baumannii*.

Natural transformation relies on the concerted expression of genes involved in DNA capture, internalization and recombination in the chromosome. DNA capture is mediated by extracellular filaments known as type IV pili (13). The Type IV pilus is a dynamic nanomachine that undergoes cycles of extension and retraction energized by at least two respective ATPase, PilB and PilT (14). Current models of Type IV pilus assembly propose that a complex of minor pilins (PilX, PilW, FimT, FimU, PilV) primes the assembly of the major pilin PilA to form a thin flexible filament (15–17). Pilus extension, capture of extracellular DNA, followed by retraction, bring DNA to the cytoplasmic space where it is bound by the ComEA receptor (13). Then DNA is converted into single-stranded DNA and translocated to the cytoplasm through the ComEC channel (18) and rapidly protected by DprA which recruits RecA for homologous recombination (19). Integration of DNA, facilitated by the ComM helicase and YraN nuclease (20), followed by replication, results in genetic transformation (21, 22).

The conditions and genetic control allowing the concerted expression of the genes involved in natural transformation – often referred to as competence – vary between species (2). For example, *Legionella pneumophila* becomes transiently competent for natural transformation in the early stationary phase when the post-transcriptional repression of transformation genes (Type IV minor pilins, comEA, comFC) by the RocC/RocR chaperone sRNA-pair is relieved (23). In *Vibrio cholerae*, chitin induces a transformation regulon, including type IV pilus genes, under control of the transcriptional activator TfoX (24). In the cyanobacterium *Synechococcus elongatus*, natural transformation is strictly regulated by circadian clock with Type IV pilus machinery produced in the morning, and natural transformation occurring at the beginning of the dark phase (evening phase) when the other genes required for transformation (including *dprA*) are expressed (25). In *Acinetobacter baumannii*, the inducing signals are undefined. Transformation protocols typically involve growth on agar or agarose-based solid medium compatible with Type IV pilus-mediated motility (7, 9, 26–30). Although the signals remained elusive, natural transformation was found to occur at the beginning of the growth phase and linked to the expression of type IV pilus genes (28).

Here, we explored the regulation of natural transformation in *Acinetobacter baumannii* by experimental conditions. Using a selection-free luminescence-based assay for natural transformation we screened a wide range of conditions, ultimately identifying a trigger of natural transformation. Genetics and molecular modeling indicate that calcium ions induce transformation by both signaling expression of the Type IV pilus and stabilization of the pilus assembly. The identified conditions allow the natural transformation of *A. baumannii* with high efficiency in liquid medium, opening the way for further investigations into the dynamics of transformation but also facilitating genetic engineering of this opportunistic pathogen.

## Results

### Mechanical sensing is not involved in induction of natural transformation

Natural transformation of *A. baumannii* is routinely done on medium solidified with agar or agarose (7, 9, 26–30). However, we and others have experienced inconsistent results that could be linked to the source of the agar or agarose used to solidify the medium. To formally establish this, we tested five references of agarose powders from four suppliers. Used at the same concentration (15 g/L) all agarose powders have identical solidifying power. We tested natural transformation of strain AB5075 using the NanoLuc (Nluc) transformation assay previously reported (10). Briefly, a suspension of AB5075 is spotted on the agarose plated together with a plasmid DNA that cannot replicate in *A. baumannii* but that carries the *nluc* gene flanked by 2 kb-long regions homologous to the chromosome. Import and recombination of this sequence leads to expression of Nluc in the transformants which can be detected by measuring luminescence with the Nluc substrate furimazine. We previously established that the luminescence signal is linearly correlated with the number of transformants over eight orders of magnitude (R^2^=0.99). Using this assay we found that only two of the tested agarose could generate transformants with RFU up to 10^6^ (Fig. 1A), corresponding to transformation frequencies of 10^-3^-10^-4^. Three other agarose gave luminescence signal barely above the value of the control without DNA (2×10^2^ RFU). This suggested that either the perception of a solid surface is required to induce transformation but some agarose contain an inhibitor, or the perception of a solid surface is dispensable for induction but some agarose contain an inducer of transformation. We tested this latter hypothesis by extracting any water-soluble compound from the unmelt agarose powder of the Euromedex D3 agarose that is permissive for transformation. A D3 agarose soluble extract (ASE) in liquid tryptone medium strongly induces transformation of strain AB5075, up to 10^7^ RLU (transformation frequency of 10^-2^), a 10,000-fold increase relative to a no-agarose liquid tryptone medium (Fig. 1B). As expected, no transformation was observed in an AB5075 *comEC* mutant. This shows that the D3 ASE contains a water-soluble potent inducer of natural transformation, aguing against a mechanical triggering of natural transformation.

**Figure 1.**
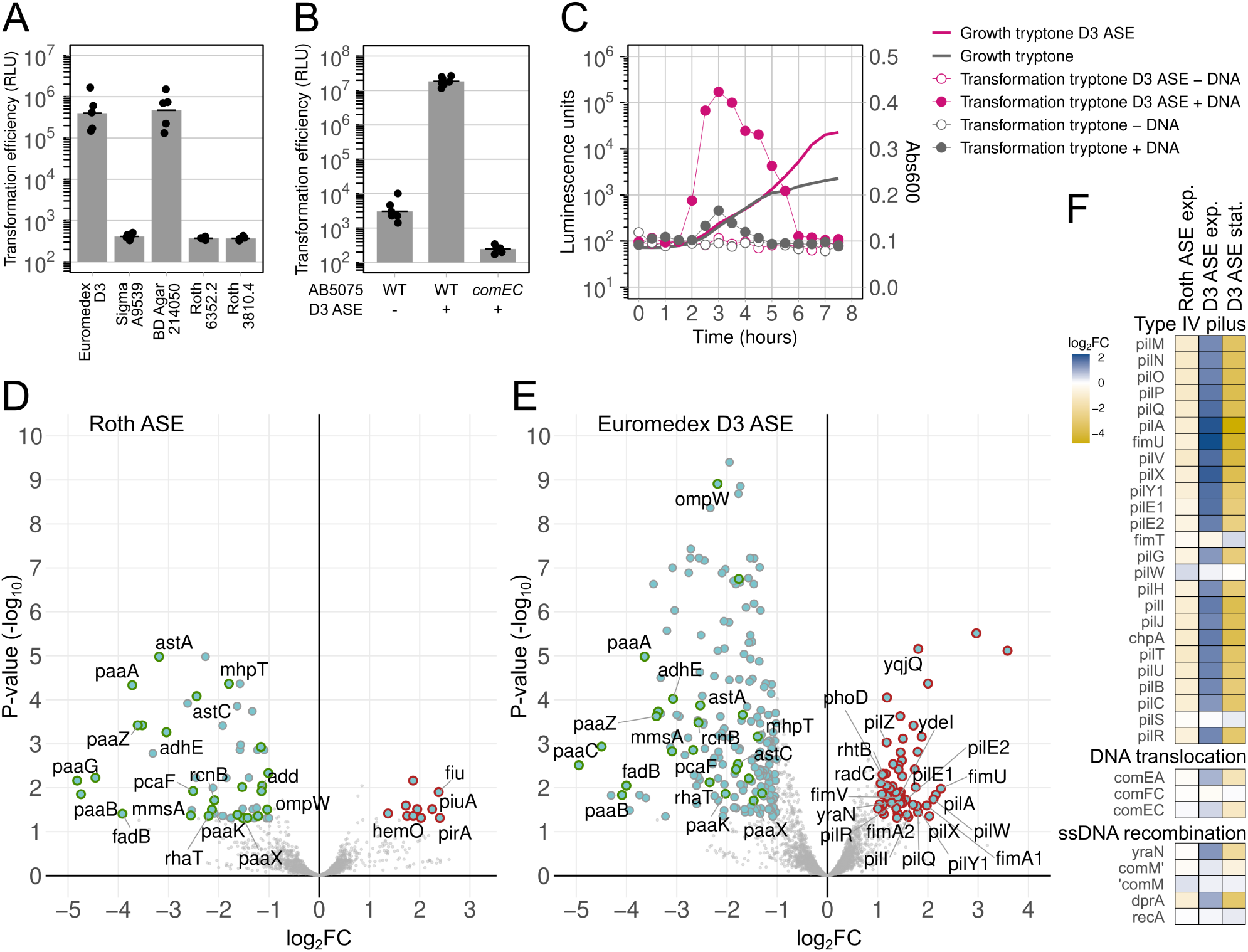
A water-soluble molecule induces a growth phase-dependent regulon of natural transformation. A. Efficiencies of natural transformation of *A. baumannii* strain AB5075 in tryptone medium solidified by different agaroses. Transformation is determined by incubating with a DNA fragment containing the Nluc gene flanked by 2 kb-long sequence of homology with the chromosome on each side. Transformants are detected by luminescence units (LU) produced when adding the substrate furimazine, and expressed relative to the optical density of the culture (RLU). The detection limit, and background luminescence is 10^2^. B. Efficiencies of natural transformation of strain AB5075 in liquid tryptone medium or tryptone agarose soluble extract (ASE) using the powder of the Euromedex D3 agarose. The method is identical as in panel A. C. Growth and natural transformation of strain AB5075 in tryptone medium or tryptone agarose soluble extract (ASE) using the powder of the Euromedex D3 agarose. Growth of the culture is monitored by reading the absorbance at 600 nm (Abs_600_). Transformation efficiency is tested at 30 minutes interval by addition of transforming DNA for 30 minutes followed by the addition of DNAse I. The culture is then further incubated. All samples were analyzed for luminescence signal, indicating effective transformation, at the end of the kinetic. D and E. RNAseq analysis of strain AB5075 grown for 3 hours in tryptone and tryptone using the non-transforming Roth agarose (D) or the transforming Euromedex D3 agarose (E). In both panel D and E, differentially-expressed genes (log_2_FC<-1 or log_2_FC>1 and P-value<0.05) are shown in blue. Genes down-regulated in both conditions are highlighted in green. Genes specifically upregulated by each condition are highlighted in deep red. F. Variation in expression of selected genes, required for natural transformation, in Roth ASE, D3 ASE and in D3 ASE at 8 hours (stationary phase) compared to D3 ASE at 3 hours (exponential phase). In AB5075, the *comM* gene is disrupted by the insertion of an AbaR element, forming two pseudogenes (*comM*’ and ‘*comM*).

### A soluble molecule induces a growth phase-dependent regulon of natural transformation

Natural transformation in *A. baumannii* was previously found regulated by the growth phase, correlating with the expression of Type IV pilus genes (28). To determine how the ASE stimulates transformation, we tested the transformability of bacteria over the growth curve. We confirmed previous results, showing that transformation peaks at 3 hours post-inoculation, corresponding to the exponential growth phase (Fig. 1C). Transformation stimulated by ASE followed the same kinetic, peaking at 3 hours, with a wider transformation window but also falling under the detection limit in the stationary phase at 7 hours post-inoculation (Fig. 1C). We then tested if increased transformation was tied to type IV pilus induction. To do so, we compared the effect of ASE from inducing (D3) and non-inducing agarose (Roth) on the global transcriptome using RNAseq profiling at 3 hours post-inoculation. Each of the non-inducing (Roth) (Fig. 2D) and inducing (D3) agarose (Fig. 2E) downregulate specific set of genes (blue dots). Yet, they also downregulate the expression of the same genes (green dots) including the phenylacetic acid catabolic pathway (*paaA*, *paaB*, *paaC*, *paaX*, *paaK*, *paaZ*). This clearly shows that adding agarose to culture medium is not without consequences on gene expression. Most importantly, while the Roth ASE induces only a dozen genes, the D3 ASE induces the upregulation of nearly all type IV pilus genes, several of which are organized in operons (*pilMONPQ*, *pilA*, *fimU-pilVWXY1-pilE1-pilE2*, *pilGHIL-chpA*, *pilBCD*), but also transformation-specific genes such as *yraN*, *dprA* or *comEA* (Fig. 1E and Fig. 1F). Notably, all of these genes are downregulated in the stationary phase (D3 ASE, 7 hours) relative to the exponential phase (D3 ASE, 3 hours) (Fig. 1F). Hence, a water-soluble compound present in agarose preparations induces a growth-phase regulated transformation regulon.

**Figure 2.**
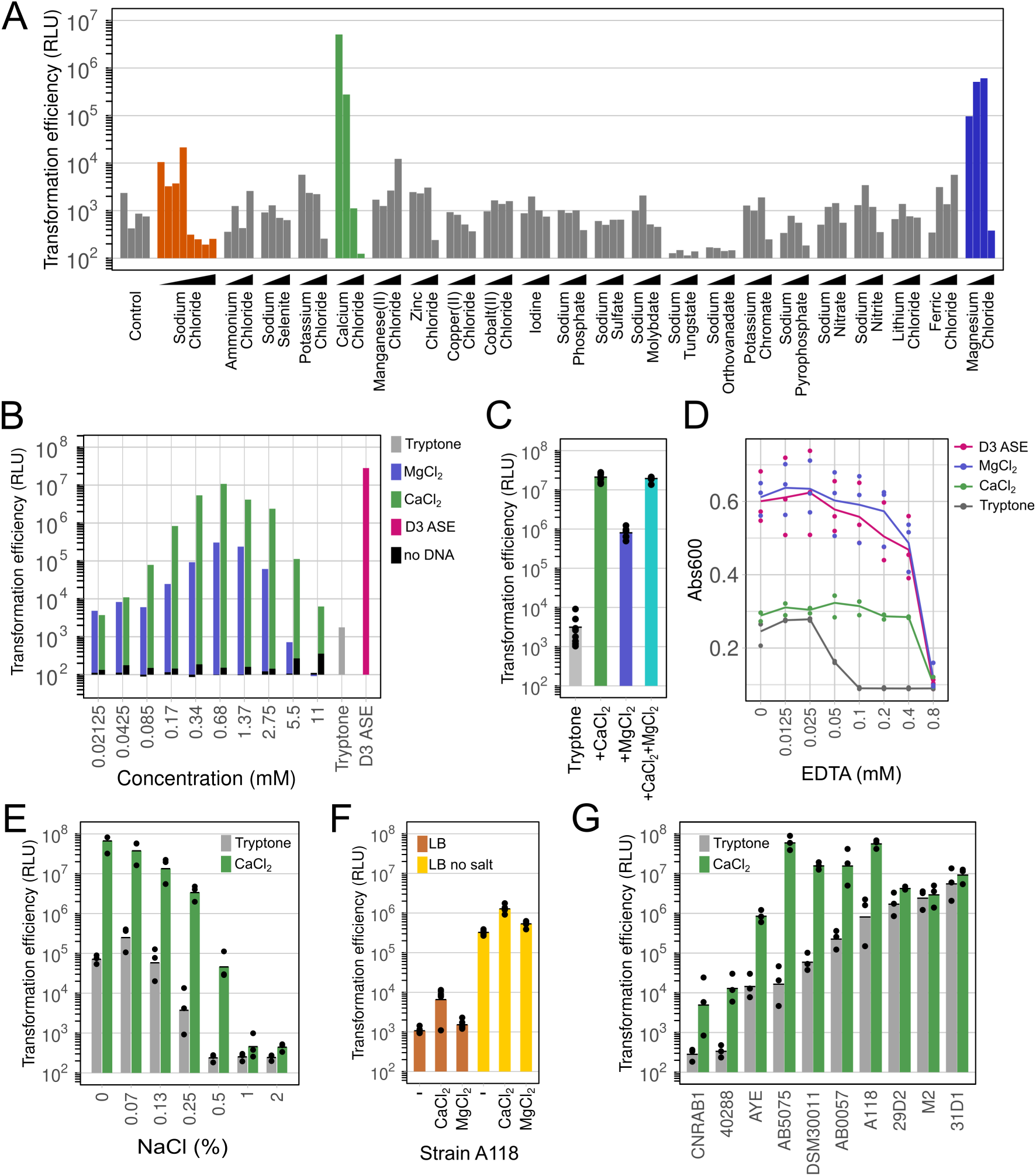
Induction of natural transformation by calcium and magnesium is antagonized by sodium chloride. A. Natural transformation of strain AB5075 in tryptone medium supplemented with the dried content of the Biolog PM-PM5 plate. B. Natural transformation in tryptone medium supplemented with increasing concentration of CaCl_2_ or MgCl_2_. C. Natural transformation in tryptone medium supplemented with CaCl_2_ (0.68 mM) or MgCl_2_ (0.68 mM) or both. Bars represent the average of technical replicates. D. Effect on growth of EDTA on tryptone medium and D3 ASE, measured at 16 hours post-inoculation by absorbance reading at 600 nm. Data are presented as average values of three independent experiments. E. Natural transformation in tryptone medium and tryptone supplemented with CaCl_2_ (0.68 mM) at an increasing concentration of NaCl. Bars represent the geometric average of three independent experiments. F. Effect of CaCl_2_ and MgCl_2_ on natural transformation of A118 in LB and LB no salt. Bars represent average of five technical replicates. G. Natural transformation of a panel of *A. baumannii* and *A. nosocomialis* (M2) strains in tryptone medium alone or supplemented with CaCl_2_ (0.68 mM). Bars represent the geometric average of three independent experiments.

### Screen identifies Ca and Mg as inducers and NaCl as repressor

RNAseq profiling revealed no upregulated genes involved in metabolism, suggesting that the transformation inducer is not metabolizable. Chemical preparations are often contaminated by ions and salt. We thus screened the Biolog PM-PM5 plates containing four undisclosed concentrations of ions for induction of natural transformation. This revealed strong induction by CaCl_2_ and to a lesser extent by MgCl_2_ (Fig. 2A). Addition of CaCl_2_ or MgCl_2_ to the tryptone medium strongly stimulated natural transformation, in a dose-dependent manner (Fig. 2B). CaCl_2_ is a stronger inducer, matching the induction by the D3 ASE (Fig. 2B), yet CaCl_2_ and MgCl_2_ showed the same dose-response effect, with maximal induction at 0.68 mM, suggesting the same mechanism of action. Indeed, the effects of CaCl_2_ and MgCl_2_ are not additive (Fig. 2C). Although CaCl_2_ is a stronger inducer than MgCl_2_, MgCl_2_ shows stimulation of growth like observed for D3 ASE in the stationary phase (Fig. 2D and Fig. 1C). This suggested that D3 ASE contained Ca^2+^ ions stimulating transformation as well as Mg^2+^ ions stimulating growth. To test this hypothesis, we evaluated the effect on growth of adding the Ca^2+^/Mg^2+^ chelator EDTA to the medium. EDTA at 0.1 mM abolished growth indicating that the tryptone medium readily contains trace amounts of Ca^2+^ (less than 0.1 mM, given the 1:1 stoichiometry of EDTA:cation). Adding CaCl_2_ at 0.68 mM did not change the growth yield (same final absorbance), but growth is expectedly inhibited at EDTA concentration of at least 0.8 mM. D3 ASE improved growth yield, identically to adding MgCl_2_ at 0.68 mM, and for both an EDTA concentration of 0.8 mM is needed to prevent growth. Thus, given the increased growth of D3 ASE and its effect on transformation similar to that of adding CaCl_2_, we conclude that D3 ASE contains both Ca^2+^ and Mg^2+^ ions, in undetermined concentrations, but likely greater than 0.68 mM.

Screening for the effect of ions on transformation, we also observed reduced transformation with increasing amounts of NaCl (Fig. 2A). This was confirmed in tryptone medium as well as when tryptone supplemented with CaCl_2_ (Fig. 2E). NaCl at concentration greater than 0.07% inhibits transformation and completely abolishes it at 1%, which is the typical concentration of the commonly used lysogeny broth (LB medium). This is consistent with previous results obtained with strain A118 (29). In LB medium, CaCl_2_ was previously reported to only moderately induce transformation in strain A118 (31). In this strain, we confirmed that CaCl_2_ has a modest effect in standard LB, and also found no effect of MgCl_2_ (Fig. 2F). The results suggest that LB might readily contain enough Mg^2+^ and a sub-optimal amount of Ca^2+^ to induce transformation but that the induction is counteracted by inhibition by NaCl. Consistent with this hypothesis, LB prepared without NaCl (LB no salt) produces a 500-fold increase in transformation, with a further increase when supplemented with CaCl_2_, while MgCl_2_ had no additional effect (Fig. 2E).

Having identified the optimal conditions for natural transformation, we tested these on a panel of strains, including commonly used laboratory strains. Transformation medium (CaCl_2_ 0.68 mM, no NaCl) improved transformation of most strains, including CNRAB1 and 40288, which were non-transformable in the absence of supplementation (Fig. 2G). Addition of CaCl_2_ also further increased transformation of the highly transformable strain A118, but did not change the transformation of the M2, 29D2 and 31D1 strains which readily showed high transformation levels in the absence of CaCl_2_ supplementation.

In conclusion, CaCl_2_ and to a lesser extent MgCl_2_, induce natural transformation. Both divalent cations are the likely contaminants contained in the transformation-inducing agarose. Adding CaCl_2_ to a tryptone medium without sodium chloride allows *A. baumannii* to undergo natural transformation in liquid medium, further demonstrating that surface sensing is dispensable for this type IV pilus-mediated process.

### Ca^2+^ and Mg^2+^ are equivalent in inducing expression of transformation genes, independently of T4P activity

In order to understand the mechanism by which CaCl_2_ and MgCl_2_ induce transformation, we analyzed the transcriptome of AB5075 exposed to CaCl_2_ or MgCl_2_ in the exponential growth phase. Consistent with D3 ASE containing calcium/magnesium salts, CaCl_2_ and MgCl_2_ induce the upregulation of the type IV pilus genes and other natural transformation genes (*comEA*, *dprA*) (Fig. 3A). The transcriptional responses to CaCl_2_ and MgCl_2_ are strikingly similar, indicating that the two metal ions are sensed by a common signaling pathway. With the *pilA* gene being upregulated and a central component of type IV pilus regulation and activity, we followed its expression using a transcriptional fusion of the full *pilA* gene to *gfp* (*pilA*::*gfp*) in a Δ*pilA* mutant. In agreement with the RNAseq data, *pilA* is induced to the same extent by CaCl_2_ and MgCl_2_ and is expressed during the exponential phase and then repressed in the stationary phase (Fig. 3B). The inhibition of transformation by NaCl suggested that it counteracts the Ca^2+^-induced expression of the type IV pilus. Indeed, in the presence of calcium, *pilA* expression is repressed by increasing concentration of NaCl, indicating that inhibition of natural transformation is the consequence of a transcriptional response to NaCl (Fig. 3C).

**Figure 3.**
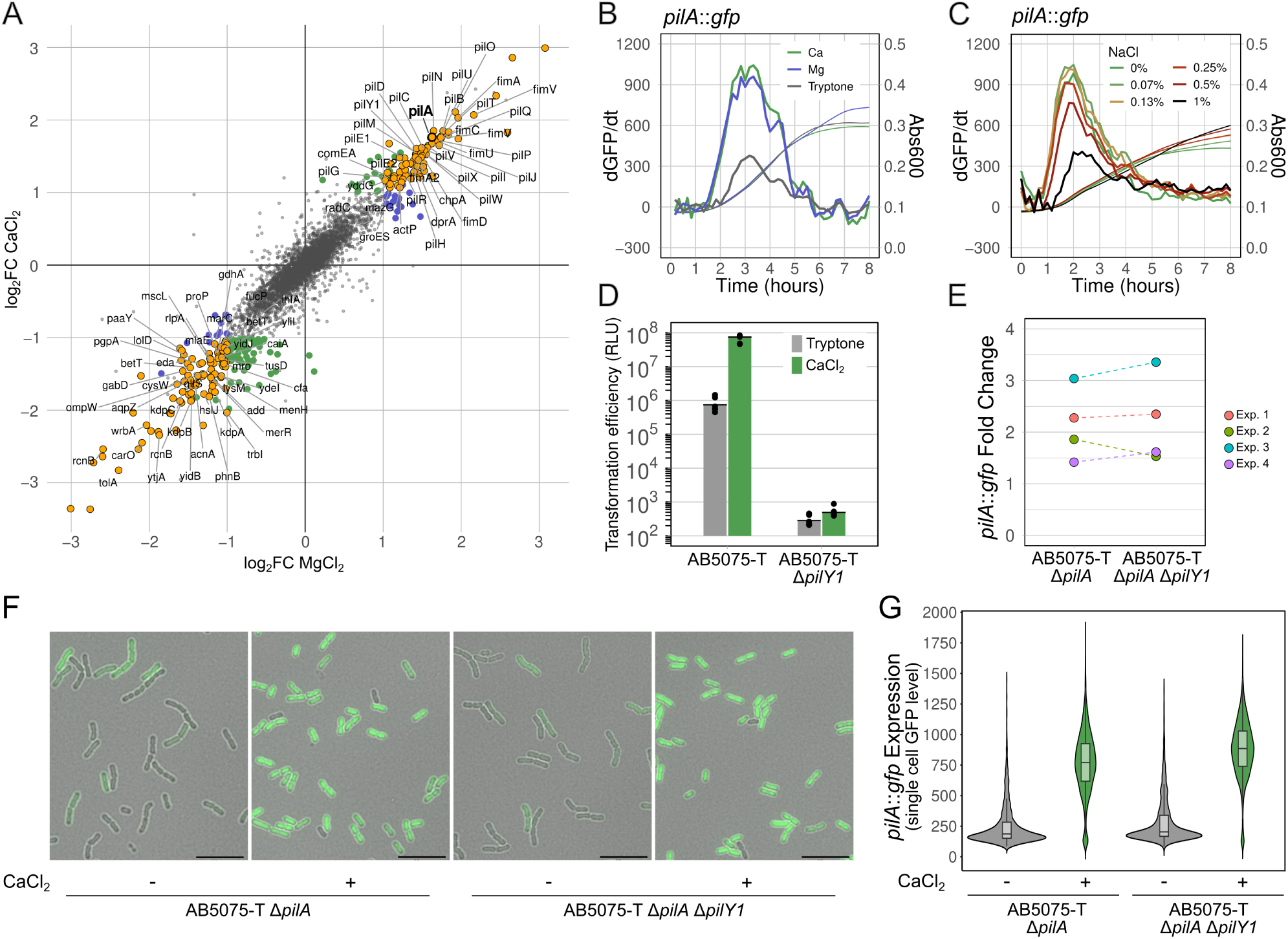
Calcium and magnesium induce a type IV pilus and transformation regulon. A. RNAseq analysis of AB5075 grown for 3 hours in tryptone medium or and tryptone medium supplemented with CaCl_2_ or MgCl_2_. Differentially-expressed genes (log2FC<-1 or log2FC>1 and P-value<0.05) in both CaCl_2_ and MgCl_2_ conditions are shown in yellow. Genes specifically differentially-regulated by CaCl_2_ and MgCl_2_ are highlighted in green and blue, respectively. B. Expression of a transcriptional fusion of the *pilA* gene to *gfp* in AB5075-T Δ*pilA*. Bacterial growth (Abs_600_) and change in *pilA*::*gfp* expression relative to cell density (dGFP/dt, arbitrary units) were measured every 10 min over an 8-h period. CaCl_2_ and MgCl_2_ were added to tryptone medium at 0.68 mM. C. Same as B, with increasing concentrations of CaCl_2_ in tryptone medium. D. The PilY1 minor pilin is essential for natural transformation in strain AB5075. Transformation conditions are identical to those of Figure 2. Presented data are an average of three independent experiments. E. Fold-change in *pilA*::*gfp* expression caused by CaCl_2_ at 0.68 mM in AB5057-T Δ*pilA* and AB5075-T Δ*pilA* Δ*pilY1*. Expression kinetics were recorded as in panel B, and fold-change represents the ratio of the maximal dGFP/dt of the CaCl_2_ condition relative to tryptone alone. Individual results from four independent experiments are shown. F. Fluorescence microscopy *pilA*::*gfp* expression in AB5057-T Δ*pilA* and AB5075-T Δ*pilA* Δ*pilY1* with or without CaCl_2_. Scale bars (bottom right corner) represent 10μm. G. Single cell expression of *pilA*::*gfp* expression in AB5057-T Δ*pilA* in tryptone medium with (n=4680, median level=772) or without CaCl_2_ (n=2449, median level=187), and AB5075-T Δ*pilA* Δ*pilY1* with (n=4373, median level=885) or without CaCl_2_ (n=2848, median level=203). Results are from two independent experiments.

The coordinated expression of the type IV pilus and the genes required for DNA import/recombination is indicative of a mechanism that equally senses Ca ^2+^ and Mg^2+^ ions and triggers a transcriptional response. It is not currently known how the *comEA* and *dprA* genes are regulated in *A. baumannii*, but regulation of type IV pilus in *A. baumannii* has some similarities with the better-known regulation in *P. aeruginosa*. The chemosensory systems Pil-Chp (32), could have been a candidate sensor/regulator, yet in *A. baumannii*, the absence of *chpA*, *pilG*, or *pilH* in transformation conditions (D3 agarose) did not change the *pilA* transcript levels (28). Expression of the major pilin PilA is under control of the two component PilRS system which can increase *pilA* transcription in response to low levels of periplasmic PilA caused by its mobilization in the extended pilus (33). In addition, in *P. aeruginosa* the minor pilin PilY1 was found to act as a Ca^2+^-dependent regulator of bacterial surface motility, favoring cycles of pilus extension/retraction (34). Hence, we hypothesized that PilY1 could sense Ca^2+^, activates pilus dynamics, which would in turn trigger *pilA* transcriptional activation through the PilRS system. To test this hypothesis, we deleted the *pilY1* gene, which, as shown for several species including *Acinetobacter baylyi* should cause complete loss of pilus assembly (34–36). As expected, the Δ*pilY1* mutant is completely defective for natural transformation (Fig. 3D). However, the loss of PilY1 did not alter the transcriptional activation of *pilA* in response to CaCl_2_ as determined at the populational level (Fig. 3E) and at the single cell level (Fig. 3F and 3G). Thus, transcriptional activation of *pilA* occurs independently of type IV pilus dynamics. In conclusion, Ca^2+^ and Mg^2+^ ions alike induce a transcriptional response mainly increasing the expression of type IV pilus and transformation genes. Interestingly, although CaCl_2_ and MgCl_2_ induce transcriptional activation of the same magnitude, CaCl_2_ can increase transformation 10- to 20-times more than MgCl_2_ (Fig. 2B and Fig. 2C), suggesting that its transformation-stimulating effect is not solely due to increased pilus expression.

### Calcium stabilizes the minor pilin complex

In *P. aeruginosa*, recent structural analyzes show that PilY1 forms a champagne cork like-structure that plugs PilQ secretin and primes the minor pilins facilitating Type IV pilus assembly and dynamics (16). Ca^2+^ binding to the conserved calcium binding loop of PilY1 is proposed to inhibit PilT-mediated pilus retraction, while in the absence of calcium pilus retraction is favored (34). We hypothesize that in *A. baumannii*, Ca^2+^ could also have a structural effect on the PilY1-minor pilin complex, directly influencing the dynamics of the pilus and, consequently, natural transformation. We first modeled PilY1 in complex with the minor pilins PilX, PilV, PilW, FimU, FimT, PilE1, PilE2 and PilA major protein using Alphafold3. Modeling of the PilY1-PilA-minor pilin complex produced a top rank model, with metrics indicating a good confidence for the overall topology (pTM of 0.72) and for inter-protein interface (ipTM of 0.73) (Fig. 4A). The modeled structure of PilY1 exhibits the characteristic architecture described in the crystal structure of PilY1 of *P. aeruginosa* (34), with an N-terminal globular head domain and a C-terminal beta-propeller fold (Fig. 4A). Importantly, the model shows that the beta-propeller fold makes direct contact with the minor pilin complex, in agreement with previously proposed pilus assembly models, based on genetic and biochemical evidence in *P. aeruginosa* (16, 37) and on cryo-electron tomography and protein interaction networks of the *M. xanthus* pilus (17). PilX is positioned directly below the C-terminal domain of PilY1 and appears to be the only direct contact with PilY1. The others minor pilin (PilV, PilW, FimT, FimU, PilE1, PilE2) and the major pilin PilA show no direct contact with PilY1 but organize into a compact helical bundle positioned beneath or beside PilX (Fig. 4A). These observations suggest that PilX may act as a structural bridge between the minor pilin complex and PilY1, potentially serving as a central element of type IV pilus initiation complex. To locate the calcium binding site on PilY1, we next modeled PilY1 in complex with PilX and performed a Ca ^2+^ docking using Alphafold3. The best model exhibits a high ranking confidence of 0.8, a pTM of 0.76, an ipTM of 0.81. Two Ca^2+^-binding motifs are predicted. One is located in the N-terminal globular head domain and has no known function, and was not present in the crystal structure of the *P. aeruginosa* PilY1 (34). The other one is located in the beta-propeller and correspond to the one required for twitching motility in *P. aeruginosa* (34) (Fig. 4B). Given that the only direct interaction of PilY1 with PilX involves the beta-propeller fold containing the calcium binding site, we next tested the role of Ca ^2+^ binding to the PilX-PilY1 interaction. To do so, we performed a molecular mechanics, generalized Born surface area calculation (MMGBSA) in presence or absence of Ca^2+^ and estimated the binding free energy (ΔG_bind) (Table1). This approach assesses how Ca^2+^ influences the stability and interaction strength between PilY1 and PilX. To ensure robustness and reproducibility, we performed two independent simulations, each on 5 structural models, including the best model presented in (Fig. 4B). In the absence of divalent cation, ΔG_bind values ranged between −195 and −210 Kcal/mol depending on the model, with negative values indicating stable interaction between PilY1 and PilX. In the presence of Ca^2+^, ΔG_bind values ranged between −212 and −230 Kcal/mol corresponding to a ΔΔG values of −18,77 Kcal/mol compared to the condition without Ca^2+^. This decrease of binding free energy value clearly supports that Ca^2+^ enhances the stability of PilY1-PilX complex. In contrast, in the presence of a Mg^2+^ ion, ΔG_bind values ranged between −188 and −207 Kcal/mol with a ΔΔG values of –2,34 Kcal/mol compared to the condition without ion. This small change in binding free energy indicates that the Mg^2+^ has a moderate role in stabilizing PilY1-PilX complex. To validate these results, we also performed the more computationally intensive molecular mechanics Poisson-Boltzmann surface area (MMPBSA) calculation. Consistent with the MMGBSA results, the MMPBSA analysis confirmed that Ca^2+^ binding strengthens the PilY1-PilX interaction (Table S3). To further investigate the role of this calcium binding loop in natural transformation, we introduced in PilY1 the substitution D947A predicted to impair Ca^2+^ coordination without altering the overall structure of PilY1 (Fig. 4C) (34, 38). In agreement with predictions, transformation efficiency is drastically reduced in the D947A substitution compared to the wild type (Fig. 4D). Taken together, the results indicate that Ca^2+^ helps stabilize the PilY1-PilX complex. Ca^2+^ binding, more than Mg^2+^, strengthens the interaction between PilY1 and PilX, which in turn promotes the recruitment of the other minor pilins and favors type IV pilus assembly.

**Table 1:** Binding free energy of PilY1-PilX. Binding free energy of PilY1-PilX (ΔG_bind) in the absence (control) or presence of Ca^2+^/Mg^2+^. Energy was calculated using MMGBSA (CHARMM36, GB8). ΔG_bind are shown for both replicate (N1,N2) of the mean of 5 independent structural models and the best model. ΔΔG values is the binding free energy difference between the control and the others conditions. A decrease of ΔΔG reflects a higher complex stability.

| Condition | replicate | $\Delta G_{\text{bind}}$<br>(best model) | Mean $\Delta G_{\text{bind}}$<br>(5 models) | $\Delta\Delta G$<br>(best model) | $\Delta\Delta G$<br>(5 models) |
| --- | --- | --- | --- | --- | --- |
| No ion | N1 | -212.69 | -194.52 |  |  |
|  | N2 | -227.06 | -209.74 |  |  |
|  | Mean | -219.88 | -202.13 | 0 | 0 |
| Calcium | N1 | -246.02 | -211.53 |  |  |
|  | N2 | -221.18 | -230.27 |  |  |
|  | Mean | -233.6 | -220.9 | -13.72 | -18.77 |
| Magnesium | N1 | -206.99 | -183.13 |  |  |
|  | N2 | -237.45 | -207.32 |  |  |
|  | Mean | -222.22 | -197.73 | -2.34 | 4.4 |

**Figure 4:**
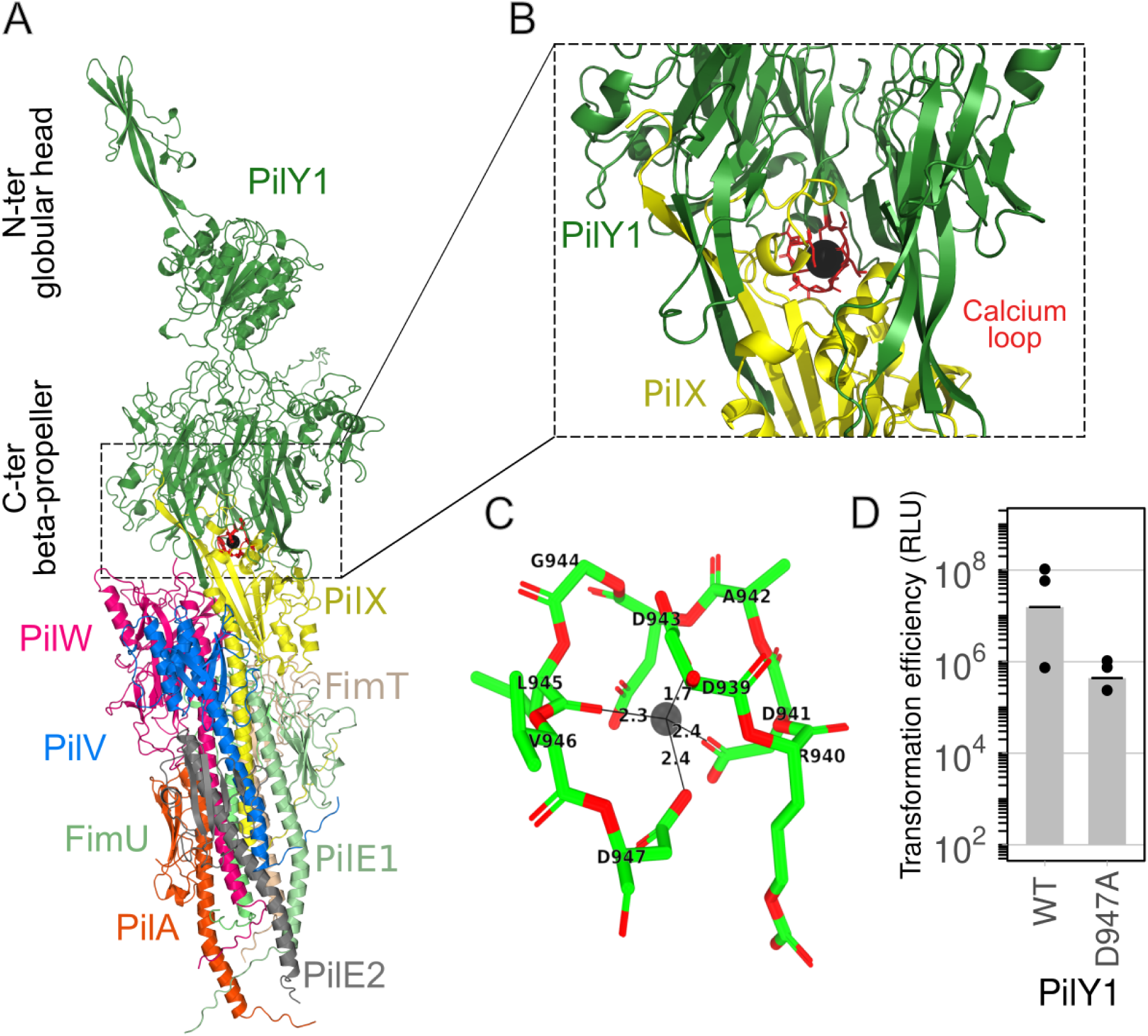
Calcium stabilizes the PilY1-minor pilin complex. A. Alphafold multimer modeling of PilY1, Type IV minor pilins and major pilin complex. Structure confidence metrics best model ipTM = 0.69, pTM = 0.71, ranking confidence = 0.73. Cartoon representation PilY1 (green), PilX (yellow), PilA (orange), PilV (blue), PilW (hotpink), FimU (lime), FimT (wheat). B. Close up on the AlphaFold model of PilY1 and PilX with calcium docking and highlighting the calcium-binding loop (red). Structure confidence metrics ipTM = 0.80, pTM = 0.76, ranking confidence = 0.81. PilY1 (green), PilX (yellow). C. Calcium binding loop modeling with relative distance in angstroms. D. Natural transformation of AB5075-T encoding the wild-type PilY1 (WT) or the PilY1 D947A substitution in the presence of CaCl_2_. Presented data are an average of three independent experiments.

## Discussion

We here report the conditions that are permissive to natural transformation in *A. baumannii* in liquid medium. Up until now, natural transformation of *A. baumannii* required the use of solid medium, and DNA uptake seemed to correlate with type IV pilus-dependent motility (26–28, 30). It could be hypothesized that surface sensing was the trigger for expression of the Type IV genes, and possibly also for other transformation genes that are needed to take up and recombine DNA. This hypothesis is supported by work in *P. aeruginosa* and *Caulobacter crescentus* showing that the surface-dependent resistance to pilus retraction is a trigger for intracellular signaling, further inducing Type IV pilus genes, virulence or chemotaxis genes (14, 39, 40). Here we found that transformation can occur in liquid medium if supplemented with submillimolar concentrations of CaCl_2_ or MgCl_2_. Under these conditions, the bacteria are not attached to a surface and cycles of extension-retraction would occur in planktonic cells, allowing the capture of extracellular DNA, likely through the binding to the minor pilin PilT (41). Activation of Type IV pilus in liquid medium is also supported by the CaCl_2_-dependent and PilA-dependent formation of large planktonic clumps of cells (Fig. S1). Activation of Type IV pilus, and processes that depend on it, can thus be achieved in the absence of surface sensing by supplementing the medium with calcium ions. Interestingly, our earlier findings suggested that CaCl_2_ and MgCl_2_ had minimal impact on transformation (9). These experiments used transformation conditions based on agarose D3, which we show here readily contains calcium and magnesium ions at levels sufficient for optimal transformation efficiency, hence explaining the lack of effect of additional calcium or magnesium ions. Another work also reported that CaCl_2_ can induce transformation in the *A. baumannii* strain A118 in LB medium but with only a 4-fold increase in transformation frequencies, a much weaker effect than the >1,000-fold increase reported here. We did confirm here the weak induction of CaCl _2_ in LB medium, and show it to be due to the presence of NaCl inhibiting transformation (Fig. 2E). Sodium chloride was indeed previously reported to inhibit transformation, causing a 100-fold decrease in transformation frequencies in A118 (29) and most other tested strains (12). Our results indicate that this is due, at least in part, to sodium chloride causing the transcriptional repression of the pilus genes (Fig. 3C).

Increases in transcript levels is a hallmark of exposure of *A. baumannii* to CaCl_2_ and MgCl_2_ alike (Fig. 3A). The transcriptional response is surprinsingly specific and coordinated, inducing all of the type IV pilus genes operons (*pilMONPQ*, *pilA*, f*imU-pilVWXY1-pilE1-pilE2*, *pilGHIL-chpA*, *pilBCD*, *pilRS*) but also transformation-specific genes such as *yraN*, *dprA* or *comEA*. This suggests that sensing of Ca^2+^ and Mg^2+^ triggers the activation of a transcriptional regulator. A possible candidate would have been the chemosensory system Pil-chp, integrating sensing by solute binding protein, yet the known response regulators of the Pil-chp system (PilG, PilH) have no role in gene expression, and most importantly, deletion of *chpA, pilG,* or *pilH* did not influence *pilA* expression (28). Another candidate was the main two-component system of the Type IV pilus system, PilRS, possibly acting through the sensing of Ca^2+^ by the type IV pilus adhesin PilY1. PilS directly senses the decrease in periplasmic PilA levels and auto-phosphorylates itself. Transfers of the phosphate to PilR activates the transcription of *pilA* (28) and, according to work on *P. aeruginosa,* also of other Type IV pilus operons (42). Ca^2+^ sensing by PilY1 could have stimulated PilA incorporation into the pilus, depleting periplasmic PilA, thus inducing a type IV pilus regulon. However, in a *pilY1* mutant that cannot sense Ca^2+^ and also cannot assemble PilA into a functional pilus, CaCl_2_ still induces *pilA* expression, indicating that it acts irrespective of the periplasmic levels of PilA sensed by PilRS (Fig 3E). The molecular sensor that can specifically interact with calcium and magnesium ions produce increased levels of mRNA of Type IV pilus genes remains unknown, and may be distinct from classical two-component system involving a transcriptional regulator. One possibility is that the transcriptional response reflects a post transcriptional mechanism. Indeed, the Ca Cl_2_ effect on *pilA* transcription appears to be moderate (∼3-fold increase in transcript) compared to the phenotypic effect (>1,000-fold increase in transformation rate). A similar situation exists in the growth phase variation of PilA protein expression which seems to exceed the 10-fold variation in its mRNA level (28). This may be the sign of a post-transcriptional regulation in which the increase in mRNA levels is not due to transcription but rather to the mRNAs being actively translated, thereby increase their stability and steady-state levels. This would be reminiscent of the control of expression of the minor pilins and the DNA uptake system proteins (ComEA, ComF, ComEC, ComM) in *L. pneumophila*, which are all under the post-transcriptional repression of a multi-target non-coding RNA (*23*). Supporting this hypothesis, a new sRNA regulator named ArpA (for *Acinetobacter* repressor of pilin) was found to target the Shine-Dalgarno sequence and the first 17 codon of *pilA* mRNA, blocking its translation and impacting natural transformation and twitching motility (43). Further work is needed to test whether ArpA, or additional sRNA, can simultaneously target in the other type IV pilus operons, *yraN*, *dprA* and *comEA*. If so, the expression of these sRNA might be modulated by calcium and magnesium ions and growth phase.

Interestingly, although CaCl_2_ and MgCl_2_ seemed identical in the activation of a transformation regulon, transformation efficiencies are about 10-times higher with CaCl _2_ than with MgCl_2_. This difference may rely on the calcium-binding loop of the non-pilin adhesin PilY1 (also named PilC1 in *Neisseria gonorrhoeae).* In *P. aeruginosa* the calcium binding site of PilY1 is required for twitching motility (34) and in *N. gonorrhoeae* the PilC1 calcium-binding mutant had a partial defect in natural transformation (38). Based on genetic and biochemical evidence, PilY1 is proposed to form a complex with minor pilins PilVWX which also interacts with PilE and FimU, acting as connector with the major pilin PilA (37). A more recent model on the *M. xanthus* pilus, based on cryo-electron tomography and protein interaction networks also supports this macromolecular organization where PilY1 sits atop the minor pilin complex (44). AlphaFold predictions of PilY1 structure with PilX, PilV, PilW, FimU, FimT, PilE1, PilE2 and PilA of *A. baumannii* is consistent with the previously proposed models, with all minor pilins forming a helical bundle linking PilY1 to the major pilin PilA. Notably, PilVWX forms a complex beneath PilY1 while FimU-PilE1E2 contact the major pilin PilA. PilY1 contacts with the minor pilin complex by interacting directly with PilX through its the calcium-binding loop. Supporting this hypothesis, the PilY1-PilX interaction is more energetically stable with Ca^2+^ coordinated in the loop. Mg^2+^ may not be the primary ligand of the PilY1 calcium loop and weakly stabilize the interaction between PilX-and PilY1. In *P. aeruginosa*, it was proposed that the PilY1 C-terminal domain bound to Ca^2+^ inhibits PilT-mediated pilus retraction, while the calcium-free PilY1 does not inhibit PilT and allows pilus retraction to proceed. The higher stability of the PilY1-PilX complex when bound to Ca^2+^ support this hypothesis, with a more stable minor pilin complex preventing disassembly and pilus retraction. The interconversion between Ca^2+^-bound and Ca^2+^-free PilY1 was proposed to control the cycles of pilus extension and retraction necessary for twitching motility. This hypothesis could be extended here to natural transformation. If Ca^2+^-binding causes less frequent disassembly, this will result in overall longer residence time of the DNA receptor FimT at the tip of the Type IV pilus (41), increasing the likelihood that DNA can be bound and imported in the periplasmic space.

The induction of natural transformation by CaCl_2_ described here is fundamentally distinct from the use of CaCl_2_ to artificially induce competence in *Escherichia coli*. In *E. coli*, high concentrations of CaCl₂ (millimolar range) are used to transiently alter membrane permeability, allowing passive entry of plasmid DNA that does not efficiently recombine with the chromosome. In contrast, the data support that Ca²⁺ in *A. baumannii* functions as a biological signal, effective at submillimolar concentrations, specifically inducing the coordinated expression and activity of the natural transformation machinery to take up and recombine extracellular DNA. Calcium ions are abundant in host-associated environments (wounds, body fluids) and on medical surfaces frequently colonized by *A. baumannii*. Whether Ca²⁺ contributes to regulation of natural transformation in these settings remains to be determined. However, Ca²⁺ is also abundant and locally variable in soil and aquatic environments and may represent an ecologically meaningful cue. Yet, the environmental reservoirs of *A. baumannii* remain poorly defined and, as of today, the ecological relevance of calcium ions for this species seems difficult to assess. Future work investigating the natural habitat of this species may benefit from including measurements of calcium ion concentrations.

Importantly, independent of its ecological and clinical significance, the addition of CaCl₂ to NaCl-free growth medium enables efficient natural transformation of *A. baumannii* in liquid culture. MgCl_2_, although less effective at stimulating transformation, improves bacterial growth and thereby increases the number of transformants recovered. Based on these observations, we established a robust protocol for natural transformation of *A. baumannii*, which will greatly facilitate genome editing by homologous recombination.

## Material and Methods

### Bacterial strains, construction and growth conditions

Bacterial strains and oligonucleotides used in this study are listed in the supplementary tables S1 and S2, respectively. Genetic modifications were made in strain AB5075 (45) or its ΔAbaR11 derivative, AB5075-T (46) and are listed in Table S2. Genetic modifications were obtained using overlap extension PCR to synthesize a large chimeric DNA fragment bordered by 2-kb fragments homologous to the insertion site and as previously described (46). For strain Δ*pilY* and *pilY1*::D947A, we adapted the multiplex genome editing by natural transformation (MuGENT) developed in *V. cholerae* (47). Briefly, this method relies on the cotransformation of two or more PCR fragments by one transformable cell. One PCR fragment carrying a selectable cassette (*aacC4* for apramycin or *tetC* for tetracycline) that integrates into a neutral genomic locus (*att*Tn*7*), is co-transformed along with PCR fragments carrying the desired mutation flanked by 2-Kb homology arms. The optimal yield was obtained when co-transforming 800 ng of non-selectable fragment with 400 ng of selectable fragment. The construction of the *pilA-sfGFP* fusion to follow *pilA* expression is described in the supplementary information along with a new chromosomal expression platform. Strains were grown either in lysogeny broth (LB) or tryptone medium (5g/L Bacto Tryptone, Gibco). All experiments were performed at 37°C. Agarose soluble extract media (ASE) was prepared by adding 0.2 g of agarose D3 (Euromedex) to 10 ml of tryptone media. This suspension was vortexed for 5 min and subsequently centrifuged to sediment the insoluble agarose particles. The supernatant was collected and filtered using a 0.22 μm filter.

### Construction of a chromosomal expression platform at the attTn*7* locus

We constructed the plasmid pAC-1 that carries the *sacB*_*aac4* selection/counter-selection cassette and a strong synthetic promoter (Pst) upstream of a reporter gene ( *gfp* amplified from from pASG-1 (9)) using an overlap extension PCR into pJET1.2 following manufacturer instructions (CloneJET PCR cloning Kit, Thermofisher scientific). Strong terminators (5S ribosomal RNA and *filA* terminators) were added to obtain the pAC-2 plasmid carrying the T_5S_-*sacB*_*aacC4*_*Pst*-*gfp*-T*_filA_*cassette. Chromosomal platforms were constructed in a two-step manner using transformation of overlap extension PCR fragments as described previously (46). First, the T_5S_-*sacB*_*aacC4*_*Pst*-*gfp*-T*_filA_*cassette was inserted in the attTn7 locus by transformation of large chimeric DNA fragment carrying the T_5S_-*sacB*_*aacC4*_*Pst*-*gfp*-T*_filA_*cassette flanked by 2 kb fragments that are homologous to the attTn7 insertion site. The resulting strain was then transformed with a chimeric PCR product carrying a fragment encompassing pilA promoter region and coding sequence and recombinants to replace the *sacB*_*aacC4*_Pst sequence. Transformants were counter-selected on minimal medium (M63) with 10% sucrose.

### Natural transformation conditions for strain constructions

Strains were grown overnight in tryptone medium, followed by dilution in tryptone medium supplemented with CaCl_2_ (0.68 mM) to obtain a final OD of 0.01. One hundred µl of culture were transferred to a 96-well plate along with 200 ng of transforming DNA and grown at 37 °C for a minimum of 7 hours, followed by plating on solid selective media (LB agar plates supplemented with apramycin, 50 µg/L or tetracycline 5 µg/L).

### Populational and single cell expression of *pilA*::*gfp*

Strains were grown for 16h in tryptone media and diluted to obtain a final optical density (OD_600_; optical path 1 cm) of 0.01 in the different media tested. Cell density in cultures in 96-well plates (Abs_600_) and fluorescence were monitored every 10 minutes over 8 hours of growth on a Tecan Infinite 200 Pro. The raw fluorescence data was smoothed using a 3-point moving average window and subsequently blank corrected using non-fluorescent wild type cells as control. OD_600_ values were similarly corrected. Expression was calculated by taking the derivative of fluorescence relative to cell density at each time point (dGFP/dT/Abs_600_).

Single cell expression was obtained by fluorescence microscopy of cultures at OD _600_ of 0.01 (as for populational analysis above) grown for 3 hours in tryptone medium with or without CaCl_2_ (1 mM). Culture were placed on agar pad on glass slides and imaged with a oil-immersion 60X objective with a EVOS M7000 microscope (Thermofisher Scientific) under phase constrast and under LED illumination for detecting GFP (482/25 nm Excitation; 524/24 nm Emission). Bacterial cells were segmented with Omnipose Cellpose (v4.08) (48) and fluorescence signal was quantified with MicrobeJ (v5.10n) (49).

### Gene Expression Analysis and Expression Profiling by RNA-Seq

Pellets from bacterial cultures in various media were stabilized and lysed as previously described using the RNAsnap method (50), followed by RNA extraction as previously described (23, 51). These samples were subsequently treated with DNase I and purified. Ribosomal-depleted RNA were used to construct strand-specific (ASE condition) or unstranded (CaCl_2_, MgCl_2_ conditions) libraries, which were sequenced on a Novaseq instrument (Illumina) (GENEWIZ Germany GmbH). Raw read processing and differential expression analyses were carried out using the Curare pipeline (52), using fastp for read trimming, Bowtie2 for alignment to the AB5075-UW reference genome (GCF_031932345.1), and DESeq2 for differential gene expression analysis. Raw reads are available at NCBI under Bioproject Accession number PRJNA1372483.

### Transformation assay using luminescence

Transformation assay using luminescence were previously described (10). It is based on the uptake and recombination of the pJET.Ab-pilMNOPQ::nLuc plasmid DNA. As this plasmid is non-replicative in *A. baumannii*, DNA molecules which are internalized undergo a double recombination event allowing the insertion of nanoluc gene in the pilMNOPQ locus and subsequent expression of the NLuc luciferase enzyme. Bacteria from an overnight culture (LB no salt, 10 g/L tryptone, 5 g/L yeast extract) were diluted at an OD _600_ of 0.01 into tryptone medium supplemented with divalent cation when indicated. In a 96-well plate, 100µl were mixed with 250ng of pJET.Ab-pilMNOPQ::nLuc plasmid DNA and incubated overnight (unless indicated otherwise) at 37°C in a static incubator. The optical density of the cell suspension at 600 nm (Abs_600_) was measured on a Tecan microplate reader. Subsequently, 80 µl of cells were mixed with 20 µl of the mix consisting of Nano-Glo® Luciferase Assay Substrate diluted 1:50 in Luciferase Assay Buffer following the manufacturer instructions (Nano-Glo® Luciferase Assay System, #N1120, Promega). Following incubation at room temperature for 10 minutes, luminescence was measured in a GloMax® Navigator Microplate luminometer (Promega). Relative luminescence units (RLU) were calculated by dividing the luminescence values by the optical density values.

### Modeling and free energy analysis of AB5075 type IV pilus initiation complex

The complete structure of the type IV pilus initiation complex comprising the major pilin PilA, the minor pilins (PilX-PilV-PilW-FimU-FimT) and the adhesin PilY1 was modeled using AlphaFold3 server (53). Five independent models were generated and evaluated using the predicted TM-score (pTM), the interface predicted score (iPTM) and ranking confidence metrics provided by Alphafold. The best model with the highest combined confidence values was selected for analysis. Interaction free energy between PilY1-PilX were estimated using the poisson-boltzmann model (PB8) gmx_MMPBSA v1.6 or the generalized born model MMGBSA (GB8) with GROMACS 2023 and the CHARMM36 force field energy with a ionic strength of 150mM as described in (54, 55). The binding free energy ΔG_bind (Kcal/mol) was extracted for each condition and average across replicate minimized model.

## Supporting information

Supplementary Figure S1 − Table S1-3 − Protocol for natural transformation

## Acknowledgements

We warmly thank Adrien Ducret for assistance in microscopy image acquisition and analysis.

