## Supplementary Figure S1 − Table S1-3 − Protocol for natural transformation for "Calcium signals natural transformation in *Acinetobacter baumannii*"

This file includes

- Figure S1: $CaCl_2$ in tryptone medium causes PilA-dependent aggregation of AB5075-T.
- Table S1: Strains used in this study
- Table S2: Oligonucleotides used in this study
- Table S3: Binding free energy of PilY1-PilX using MMPBSA calculation
- Protocol for natural transformation of Acinetobacter baumannii in liquid medium

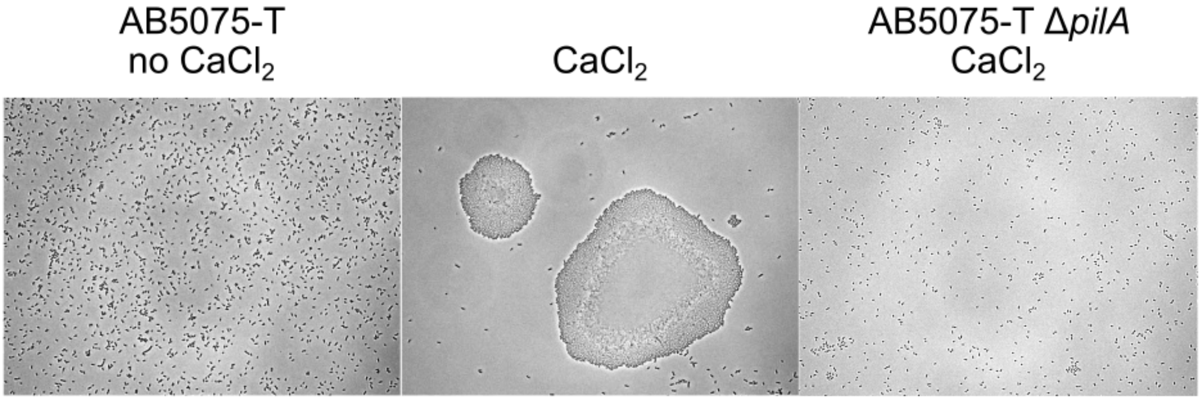


**Figure S1: $CaCl_2$ in tryptone medium causes PilA-dependent aggregation of AB5075-T.** Overnight cultures of AB5075-T and pilA mutant were directly imaged under white light illumination.

#### Table S1: Strains used in this study

| Strain name | Genotype/comment | Reference or source |
|---|---|---|
| AB5075-T | wild type | (45) |
| AB5075 | wild type | (44) |
| AB5075 *att*Tn*7::sacB_aacC4_Pst-sfGFP* | *att*Tn*7::sacB_aac4_Pst-sfGFP* | this study |
| AB5075-T *pilA-sfGFP ΔpilA* | *att*Tn*7::pilA-gfp _aac4 ΔpilA* | this study |
| AB5075-T *pilA-sfGFP ΔpilA ΔpilY1* | *att*Tn*7::pilA-gfp_aac4 ΔpilA ΔpilY1* | this study |
| AB5075-T ΔcomEC | *comEC::sacB_aacC4* | this study |
| AB5075-T Δ*pilY1* | *att*Tn*7::tetC ΔpilY* | this study |
| AB5075-T *pilY1::D947A* | *att*Tn*7::tetC pilY(*D947A) | this study |
| CNRAB1 | wild type | (7) |
| 40288 | wild type | (53) |
| AYE | wild type | (54) |
| DSM30011 | wild type | (55) |
| AB0057 | wild type | (56) |
| A118 | wild type | (57) |
| 29D2 | wild type | (58) |
| M2 | wild type | (26) |
| 31D1 | wild type | (58) |

#### Table S2: Oligonucleotides used in this study

| Genetic construction | Name | Sequence 5' to 3' | Template; Annealing site |
|---|---|---|---|
| ***Chromosomal modifications using overlap extension PCR*** | | | |
| *att*Tn*7::sacB_aacC4_Pst-gfp* | oac-6 | tggtaatggagcaacagtcg | AB5075 genomic DNA; 2kb upstream *att*Tn*7* site at the end of AB5075 *glmU* gene |
| | oac-7B | ggaacttcgaagcagctccagcctacacaatcctgacttcggctacacatctagaattg | AB5075 genomic DNA, 5S terminator-near site *att*Tn*7*, downstream AB5075 *glmS* terminator, and an adaptor |
| | oac-8B | gaactaaggaggatattcatatggaccatggcgtgatcttaggggtgataattctg | AB5075 genomic DNA, *filA* terminator-site near *att*Tn7, and an adaptor |
| | oac-9 | aaacgctagaacgagacggg | AB5075 genomic DNA; 2kb downstream *att*Tn7 site in AB5075 genomic DNA |
| | oac-4B | gattgtgtaggctggagctgcttcgaagttcctattctaaacaccccaatctcaaaagag | Annealing to pAC2, share a common sequence with oac-4 (partial terminator and adaptor) |
| | oac-5B | gccatggtccatatgaatatcctccttagttcaagtatttataaagagcgcgtttatacg | Annealing to pAC2, share a common sequence with oac-5 (partial terminator and adaptor) |
| | oac-13 | atagtgaacggcaggtatatgtgatgggtg | AB5075 *att*Tn7::*sacB-aac-Pst-gfp* genomic DNA, annealing on the *sacB* gene but downstream the 5S terminator |
| *pilA* gene and its promoter region | mlo-142 | cacccatcacatatacctgccgttcactataagcggtagttacaggagttcagc | AB5075 genomic DNA; 450nt upstream *pilA* |
| | mlo-143 | gggtaccgagctcgaattctgtttcctgtgcttatgctgcagggcaagcagaac | AB5075 genomic DNA; *pilA* stop codon |
| ***Primers for pAC-1 construction*** | | | |
| *sacB_aacC4* | mlo-86 | gattgtgtaggctggagctgcttcgaagttcctcacccatcacatatacctgccgttcac | Annealing on pMHL2 |
| | oac-1 | ccgcgggattacaggttggatgataagtc | |
| *Pst-gfp* | oac-2 | tatcatccaacctgtaatcccgcggggcgaaaatcctgtttgatggtgg | Annealing on pASG1 |
| | oac-3 | gccatggtccatatgaatatcctccttagttcttatttgtagagctcatccatgccgtgc | |
| ***Primers for pAC-2 construction*** | | | |
| Addition of terminators to the *sacB_aacC4_Pst-gfp* | oac-4 | tattctaaacaccccaatctcaaaagaggttggggtgttttttatatcacccatcacatatacctgccgttcac | annealing on pAC1, adds 5S terminator upstream *sacB* |
| | oac-5 | aagtatttataaagagcgcgtttatacgcgctctttatttatcttttatttgtagagctcatccatgccgtgc | Annealing to pAC1, adds *filA* terminator |

| | | | |
|---|---|---|---|
| cassette | | | downstream *gfp* |
| ***Primers for ΔpilY markless construction*** | | | |
| Markerless deletion of *pilY1* | 80_Mutgent_delta_pilY_F1 | CGATTGCGGTCATGGCTATTATTG | AB5075 genomic DNA; 3kb upstream *pilY1* gene |
| | 81_Mutgent_delta_pilY_R1 | CAGAGCTTGTCATCAGCCATG | AB5075 genomic DNA; upstream *pilY1* gene |
| | 82_Mutgent_delta_pilY_F2 | CAATGATATTCACATGGCTGATGACAAGCTCTGCTAATTGCCCAGTAGGGGCT | AB5075 genomic DNA; downstream *pilY1* gene |
| | 83_Mutgent_delta_pilY_R2 | CAAACTGGTCTGGCTTAGTGTATTG | AB5075 genomic DNA; 3kb downstream *pilY1* gene |
| ***Primers for pilY::D947A markless construction*** | | | |
| markless substitution of *pilY1 Aspartate 947 to Alanine* | KG29_pilY_Noscare_F1_F | GCCTCGACTTCTTTAGAAACTC | AB5075 genomic DNA; 3kb upstream *pilY1* gene |
| | KG36_PilY_DtoA_F1_R | CTCCAAAATATAGATGGGCGACTAAGCCATCAGCATCACG | AB5075 genomic DNA; in PilY1 calcium loop |
| | KG37_PilY_DtoA_F2_F | CGTGATGCTGATGGCTTAGTCGCCCATCTATATTTTGGAG | AB5075 genomic DNA; in PilY1 calcium loop |
| | KG38_PilY_DtoA_F2_R | CCGCAGATGCAGTAGCTCTAC | AB5075 genomic DNA; 3kb downstream *pilY1* gene |
| ***Primers for construction of mutgent cassette att*Tn*7::tetC*** | | | |
| *Insertion of tetC into* ***att*Tn*7* site of AB5075** | 72_K7_tc_PCR1_F1 | tggtaatggagcaacagtcgg | AB5075 genomic DNA; 3kb upstream *att*Tn*7 site* |
| | 73_K7_tc_PCR1_R1 | GATTGTGTAGGCTGGAGCTGCTTCGAAGTTCCctgacttcggctacacatctagaattg | AB5075 genomic DNA; into *att*Tn*7 site* |
| | 74_K7_tc_PCR2_F2 | GGAACTTCGAAGCAGCTCCAGCCTACACAATCCTGTCTCTTATACACATCTCAACCA | T26 |
| | 75_K7_tc_PCR2_R2 | GCCATGGTCCATATGAATATCCTCCTTAGTTCCTGTCTCTTATACACATCTCAACCCT | T26 |
| | 76_K7_tc_PCR3_F3 | GAACTAAGGAGGATATTCATATGGACCATGGCgtgatcttaggggggtgataattctg | AB5075 genomic DNA; 3kb into *att*Tn*7 site* |
| | 77_K7_tc_PCR3_R3 | aaacgctagaacgagacggg | AB5075 genomic DNA; 3kb downstream *att*Tn*7 site* |
| ***Primers for construction of mutgent cassette att*Tn*7:: att*Tn*7::pilA-gfp _aac4*** | | | |
| *Insertion of aac4 into* ***att*Tn*7* site of AB5075** | 72_K7_tc_PCR1_F1 | tggtaatggagcaacagtcgg | AB5075 *attTn7::pilA-gfp* genomic DNA; into *att*Tn*7 site* |
| | PilAGFP-P1-R2 | GCCATGGTCCATATGAATATCCTCC | AB5075 *attTn7::pilA-gfp* genomic DNA; into *att*Tn*7 site* |
| | PilAGFP-P2-F1 | GAACTAAGGAGGATATTCATATGGACCATGGCacaggttggatgataagtccc | Annealing on pMHL10; acc4 amplification |
| | PilAGFP-P2-R2 | atgtgttggcagccttttcagcacatcactCTCATGAGCTCAGCCAATCG | Annealing on pMHL10; acc4 amplification |
| | PilAGFP-P3-F1 | agtgatgtgctgaaaaggctgc | AB5075 genomic DNA; 3kb into *att*Tn*7 site* |
| | 77_K7_tc_PCR3_R3 | aaacgctagaacgagacggg | AB5075 genomic DNA; 3kb downstream *att*Tn*7 site* |

| ***Primers for amplification of existing ∆pilA fragment*** | | | |
|---|---|---|---|
| *Insertion of aac4 into* ***att*****Tn*****7 site of AB5075*** | 95_PilA_Amp_F1 | CGGCTTGGTCTGAAGTACAAAAC | AB5075 *attTn7::pilA-gfp ∆pilA* genomic DNA; into *pilA site* |
| | 94_PilA_Amp_R2 | GCTCAACCAAGTGCTGTATCATC | AB5075 *attTn7::pilA-gfp ∆pilA* genomic DNA; into *pilA site* |

**Table S3: Binding free energy of PilY1-PilX using MMPBSA calculation**

| Condition | replicate | ΔG_bind (best model) | Mean ΔG_bind (5 models) | ΔΔG (best model) | ΔΔG (5 models) |
|---|---|---|---|---|---|
| No ion | N1<br>N2 | -199,75<br>-220,37 | -178,86<br>-201,3 | | |
| | Mean | -210,06 | -190,08 | 0 | 0 |
| Calcium | N1<br>N2 | -241,21<br>-234,98 | -200,44<br>-218,76 | | |
| | Mean | -238,095 | -209,60 | -28,04 | -19,52 |
| Magnesium | N1<br>N2 | -179,31<br>-225,73 | -166,59<br>-200,83 | | |
| | Mean | -202,52 | -183,71 | 7,54 | 6,37 |

Table of binding free energy of PilY1-PilX (ΔG_bind) in the absence (control) or presence of $Ca^{2+}$/$Mg^{2+}$. Energy was calculated using MMPBSA (CHARMM36, PB8). ΔG_bind are shown for both replicate (N1,N2) of the mean of 5 independent structural models and the best model. ΔΔG values is the binding free energy difference between the control and the others conditions. A decrease of ΔΔG reflects a higher complex stability.

Horigene Team – CIRI Lyon France.
Protocol created by Jason Chirakadavil, updated Jan, 27th, 2026

**From "Calcium signals natural transformation in *Acinetobacter baumannii*"**

by Jason Baby Chirakadavil, Kelly Goldlust, Ludovic Poiré, Nicolas Gaudin, Alexandre Chassard, Adrien Camilli, Manon Bouvier, Maria-Halima Laaberki, Xavier Charpentier

### Protocol for natural transformation of *Acinetobacter baumannii* in liquid medium

#### Introduction

This is the new protocol for natural transformation in *Acinetobacter baumannii* (also tested with *Acinetobacter nosocomialis)* in liquid medium. This is possible because Ca++ and Mg++ ions induce the genes required for natural transformation, while presence of NaCl inhibits natural transformation. Once sterilized, the media components can be reconstituted and stored at RT without any loss in transformation efficiency.

Ca++ is sufficient to induce transformation. Mg++ is less efficient in inducing transformation but it stimulates growth, so combining both can slightly increase the number of transformants that can be obtained. Yet, transformation is very efficient, so Mg++ can be considered optional.

#### Materials

- LB (without NaCl)
    - Mix the following components in MilliQ water
    - 10g/L Tryptone (Thermofisher Bacto™ Tryptone. Catalog number: 211705)
    - 5g/L yeast extract (Thermofisher Gibco™ Yeast Extract: Catalog number: 211929)
    - Sterilize by autoclaving
- Tryptone media (without NaCl)
    - Mix the following components in MilliQ water
    - 5g/L Tryptone (Thermofisher Bacto™ Tryptone. Catalog number: 211705)
    - Sterilize by autoclaving
- LB agar plates with appropriate antibiotics for selection
- Transformation Inducers
    - Mix the following components in MilliQ water
    - $CaCl_2$ 680 mM solution - 1000X stock
    - $MgCl_2$ 680 mM solution - 1000X stock (optional)
    - Sterilize by filteration using a 0.2 µm filter

**Procedure**

- **Preparation of "transformation media"**

1. Dilute the $CaCl_2$ and $MgCl_2$ (optional) at 680 mM into tryptone media to get a final concentration of 680 μM (1 μl $CaCl_2$ in 1000 μl tryptone media)
2. Store solution at RT

- **Typical transformation experiment**
- **Day 1**

3. Start a preculture of *Acinetobacter baumannii* either from a plate or scrape cells directly from the glycerol stock into 2 ml of LB (without NaCl) in a 13 ml sterile culture tube.
4. Incubate overnight (16h) at 37°C with shaking (~200 rpm)

- **Day 2**

5. Check the $OD_{600nm}$ of the preculture (should be around 5-8)
6. Aliquot 100 μl of transformation media into a 1.5 ml eppendorf tube

- Alternatively, 100 ul of transformation media into a 96-well plate (if you have multiple transformations to perform)

7. Add cells from the preculture into transformation media to obtain $OD_{600nm}$ of 0.01
8. Add ~2 ng/μl transformation DNA (200 ng each tube/well) (see Note 1), include control tubes without added transformation DNA
9. Incubate at least 5 hours at 37°C without shaking

- Alternatively, incubate overnight (~16h) at 37°C without shaking (see Note 2)

10. Plate 100 μl of cells on LB plates
11. Incubate the plates overnight at 37 °C

- **Day 3**

12. Pick colonies that grew in the appropriate selection conditions

**Notes :**

1. DNA concentration of 2 ng/μl was used in our assay, however this depends on the quality of the transforming DNA. If assembled PCR products are to be used, you can add up to ~5 ng/μl.

2. The peak of transformation occurs at ~3 hrs post inoculation at 0.01 OD. Hence, you can try plating the cells after 3.5 hours, but the number of transformants might be lower.

3. It is absolutely essential that the **transformation media** contains **no NaCl**

4. The tryptone medium does not age well, growth yield can decrease over time. Avoid long-term storage (weeks).
